# Corticotropin-Releasing Factor Neurons in the Bed Nucleus of the Stria Terminalis Differentially Influence Pain Processing and Modulation in Male and Female Mice

**DOI:** 10.1101/2020.07.24.219451

**Authors:** Waylin Yu, Christina M. Stanhope, Natalia del R. Rivera Sanchez, Garrett A. Moseley, Thomas L. Kash

## Abstract

The bed nucleus of the stria terminalis (BNST) plays an emerging yet understudied role in pain. Corticotropin-releasing factor (CRF) is an important source of pain modulation in the BNST, with local pharmacological inhibition of CRF receptors conditionally impacting the sensory and affective-motivational components of pain. Knowledge on how pain dynamically engages CRF neurons in the BNST and is influenced by intra-BNST production of CRF, however, remains unknown. In the present study, we utilized *in vivo* calcium imaging to show robust and synchronized recruitment of BNST^CRF+^ neurons during acute exposure to noxious heat. Distinct patterns of recruitment were observed by sex, as the magnitude and timing of heat responsive activity in BNST^CRF+^ neurons differed for male and female mice. We then established the necessity of CRF for intact pain behaviors by genetically deleting *Crf* in the BNST, which reduced thermal and mechanical nociceptive sensitivity for both sexes, and increased paw attending in female mice, suggesting a divergent role for CRF with respect to active coping responses to pain. Together, these findings demonstrate that CRF in the BNST contributes to multiple facets of the pain experience and may play a key role in the sex-specific expression of pain-related behaviors.

## Introduction

Pain is a pervasive and well-conserved source of stress that disrupts homeostasis by driving negative sensory and emotional states (Abdallah & Geha, 2017). Corticotropin-releasing factor (CRF), a peptide known for regulating stress-related behaviors, has been hypothesized to play an impactful, albeit understudied, role in pain modulation (Wei et al., 1986; Hargreaves et al., 1987; Valentino & Foote, 1987; Song & Takemori, 1991; Kita et al., 1993; Mousa et al., 1996; Schafer et al., 1996; Lariviere & Melzack, 2000; Cui et al., 2004; Sinniger et al., 2004; Mousa et al., 2007; Zhang & Xu, 2017). Recent work has highlighted the bed nucleus of the stria terminalis (BNST) as a CRF-enriched structure that can significantly influence the affective-motivational components of pain (Rouwette et al., 2011; Minami & Ide, 2015, Minami, 2019). Bilateral lesions of the BNST suppress the aversive aspects of electrical, visceral, and somatic pain in rats (Crown et al., 2000; Deyama et al., 2007; 2008; 2009). Furthermore, CRF release is increased in the BNST following intraplanar injection of formalin, a tonic-acting inflammatory agent that exacerbates sensitivity to noxious stimuli (Ide et al., 2013), and intra-BNST administration of CRF results in conditioned place aversion in a CRF_1_ and CRF_2_ receptor-dependent manner (Ide et al., 2013). Notably, these studies reported no effect on formalin-induced nocifensive behaviors (Ide et al., 2013; Kaneko et al., 2016). By contrast, intra-BNST antagonism of CRF_2R_ reduces mechanical and colonic nociception, and CRF_1R_ antagonism attenuates stress-induced hyperalgesia (Tran et al., 2012; 2014). The discrepancies in CRF contributions to pain sensitivity reported in these studies indicate the need for additional evaluations on the conditions by which CRF signaling in the BNST can alter the sensory and affective-motivational components of pain.

To gain a better understanding of CRF signaling in the BNST and its contributions to pain, two key functional components require clarification. (1) To date, the majority of studies focusing on the BNST and pain have been limited to the impact of local CRF receptors, notably lacking insight on the sources of CRF that influence the conditional effects of CRF receptor antagonism on pain (Nijsen et al., 2005; Tran et al., 2012; 2014; Ide et al., 2013). Whether it is possible to manipulate CRF signaling in the BNST at the peptide level to produce similar outcomes remains to be determined. (2) Previous studies were performed exclusively in male subjects, despite evidence that CRF signaling in the BNST drives discrete stress responses in male and female rodents (Janitzky et al., 2014; Klampfl et al., 2016; Smithers et al., 2019). Sex differences in pain expression and treatment are well-established in humans (Paller et al., 2009; Bartley & Fillingim, 2013), but preclinical pain studies have historically excluded female subjects (Mogil et al., 2012), thus increasing the potential for neglected mechanistic insight. These uncharacterized features of CRF signaling reveal significant gaps in our understanding of how the BNST regulates pain.

In the present study, we sought to establish the BNST as a possible source of CRF for the modulation of CRF_1/2R_-dependent pain-related behaviors in male and female mice. Given the high levels of CRF peptide and receptor expression in the BNST, it is possible that local production of CRF plays an important role in the regulation of pain (Cummings et al., 1983; Sakanaka et al., 1986; Ju et al., 1989; Moga et al., 1989; Morin et al., 1999; Dabrowska et al., 2013; Daniel & Rainnie, 2016). Previous descriptions of the BNST as a sexually dimorphic structure additionally make it possible for CRF to play discrete roles in pain for male and female subjects (del Abril & Guillamon, 1987; Han & De Vries, 2003; Bangasser & Shors, 2008). To test these hypotheses, we characterized the contributions of CRF to pain processing and modulation in both sexes by using genetically targeted manipulations of the peptide in the BNST. First, we used *in vivo* calcium imaging to examine how BNST^CRF+^ neurons encode for acute exposures to noxious heat. We then assessed pain-related behaviors following genetic deletion of CRF in the BNST, allowing us to discern peptide-specific contributions to both acute and sustained pain. By tracking endogenous BNST^CRF+^ activity and manipulating peptide expression in both sexes, the present study highlights a crucial function of the BNST as a locus of CRF signaling and sex-dependent pain regulation.

## Materials and Methods

### Animals

Male and female CRF-Cre (Martin et al., 2010) and Floxed-CRF (Zhang et al., 2017) mice (N = 103 total, ≥ 6 weeks of age, age-matched for each experiment, C57BL/6J background) were bred in-house, group-housed with same-sex littermates, and maintained on a 12-h light/dark cycle (light on at 7:00, light off at 19:00) with rodent chow and water available *ad libitum*. Subjects were singly housed for imaging experiments to ensure recovery and prevent post-surgical complications. All procedures were approved by the Institutional Animal Care and Use Committee at UNC Chapel Hill and performed in accordance with the NIH Guide for the Care and Use of Laboratory Animals.

### Surgeries

Subjects were anesthetized with isoflurane (1-3%) in oxygen (1-2 l/min) and aligned on a stereotaxic frame (Kopf Instruments, Tujunga, CA). All surgeries were conducted using aseptic techniques in a sterile environment. Microinjections were performed with a 1 μl Neuros Hamilton syringe (Hamilton, Reno, NV) and a micro-infusion pump (KD Scientific, Holliston, MA) that infused virus at 100 nl/min. Viruses were administered bilaterally in the dorsal region of BNST (250 nl for all experiments; relative to bregma: ML ±0.90 mm, AP +0.23 mm, DV −4.35 mm). For experiments that required *in vivo* calcium imaging, Gradient-index (GRIN) lenses were implanted unilaterally in the right hemisphere approximately 200 μm over the dorsal BNST (relative to bregma: ML ±0.90 mm, AP 0.23 mm, DV −4.15 mm) and secured with a dental cement headcap. Baseplates were later added to stabilize the attachment of the miniature microscope (see “*Ca*^2+^ *imaging with miniature microscope*” section for details). After surgery, mice were given *ad libitum* Tylenol water or daily injections of meloxicam (5 mg/kg, subcutaneous [s.c.]) for four consecutive days, then allowed to recover for three weeks or longer before starting experiments.

### Behavioral assays

#### Hargreaves

Subjects were placed in Plexiglas boxes on an elevated glass surface and habituated to the behavioral apparatus for 30-60 minutes to reduce novelty-induced locomotion and other factors that may confound nociceptive sensitivity measures. The mid-plantar surface of each hind paw was exposed to a series of noxious heat trials that sequentially alternated between left and right paws. After each heat exposure, a 10-minute inter-trial interval was provided before the next trial. Six trials of heat exposure were conducted for all experiments except those using *in vivo* calcium imaging, which we restricted to 4 trials to prevent photobleaching. For Figures 1–2 (BNST^CRF+^ imaging experiments), beam intensity was set to 15 on the IITC Plantar Analgesia Meter (IITC Life Science, Woodland Hills, CA), producing average basal paw withdrawal latencies of approximately 10 seconds in CRF-Cre mice. For Figures 3–5 (*Crf* deletion experiments), beam intensity was set to 25 on the IITC Plantar Analgesia Meter, producing average basal paw withdrawal latencies of approximately 6 seconds in Floxed-CRF mice. Cutoff times of 20 seconds were set for each trial to prevent excessive tissue damage.

**Figure 1.**
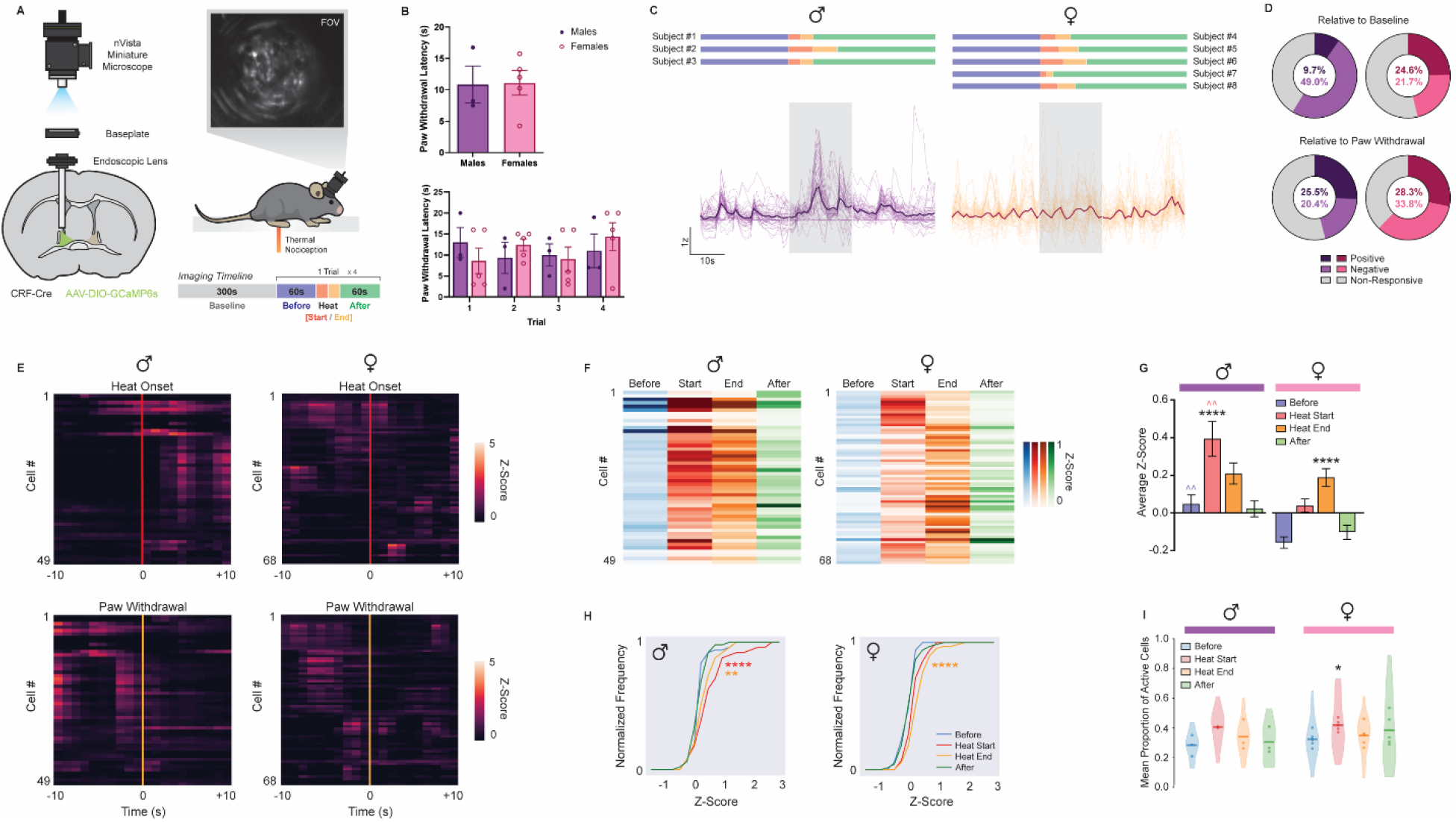
Measurement of Single-Cell Ca^2+^ Activity in BNST^CRF+^ Neurons During Pain. **(A)** Miniature microscope imaging in BNST^CRF+^ neurons. (Left) Diagram of GCaMP6s infusion and endoscopic lens / baseplate implantation in the BNST of CRF-Cre mice. (Upper-Right) Imaging field-of-view (FOV) depicting GCaMP6s expression in BNST^CRF+^ neurons. (Lower-Right) Experimental timeline with schematic of BNST^CRF+^ imaging during thermal nociceptive sensitivity testing. Epochs relative to heat exposure are color-coded: Before (blue), Heat Start (red), Heat End (orange), and After (green). **(B)** (Left) Thermal nociceptive sensitivity of male (n = 3) and female (n = 5) CRF-Cre mice, as measured by the average paw withdrawal latency (PWL) following four trials of the Hargreaves test (unpaired t-test: t(6) = 0.07921, p = 0.9394). (Right) Average PWL of male and female subjects by trial (Two-way mixed-model ANOVA with Sidak’s post hoc: no Trial × Sex interaction [F(3, 18) = 1.159, p = 0.3526] or main effect of Trial [F(2.498, 14.99) = 0.5867, p = 0.6041]; main effect of Sex [F(1, 6) = 0.0062, p < 0.0001]). **(C)** Relative fluorescence change (ΔF/F) of BNST^CRF+^ activity during noxious heat exposure. Z-scores of male (n = 49 cells / 7-30 cells per mouse) and female (n = 68 cells / 8-19 cells per mouse) subjects are represented over time by aligning traces that were averaged across trials to the onset of the heat stimulus. Individual cell activity is indicated in lighter colors, with total average cell activity in darker colors, and maximum heat duration (20 seconds) in the transparent blue block. Scale bars, × = time (10 sec), y = z-score based on relative ΔF/F (1 z-score). Imaging timelines for individual subjects are located above to represent epochs via the color-coding system described in **(A)**. **(D)** Total percentage of pain responsive cells for male (purple) and female (magenta) subjects across four trials. Cells that respond to noxious heat exposure with positive (darker), negative (lighter), and no (grey) changes in z-score were determined relative to the *Before* and *After* epochs, as demarcated by the onset of heat exposure and paw withdrawal (Wilcoxon rank-sum, p < 0.05). **(E)** Peri-event heatmap of average BNST^CRF+^ activity surrounding heat onset (red line) and paw withdrawal (orange line). **(F)** Average z-score of each neuron across epochs for male and female subjects. **(G)** Average z-score by epoch for male and female subjects (Two-way mixed-model ANOVA with Tukey’s post hoc: Sex × Epoch interaction [F(3, 345) = 5.996, p = 0.0005], main effect of Sex [F(1, 115) = 13.20, p = 0.0004] and Epoch [F(2.153, 247.7) = 26.31, p < 0.0001]). * denotes comparison of epochs for each sex, ^ denotes comparison of the same epoch between sexes. **(H)** Cumulative distribution function comparing z-score frequency for each epoch in male and female subjects (KS test: By Sex – Before [D(117) = 0.3466, p value = 0.0014], Heat Start [D(117) = 0.3850, p value = 0.0002], Heat End [D(117) = 0.1815, p value = 0.2759], After [D(117) = 0.2611, p value = 0.0334]; Comparing each epoch within males results in significance with Heat Start and Heat End over Before and After [Before vs. Heat Start, D(98) = 0.4694, p value = 2.183e-05; Before vs. Heat End, D(98) = 0.4286, p value = 0.0001; Heat Start vs. After, D(98) = 0.4082, p value = 0.0003; Heat End vs. After, D(98) = 0.3265, p value = 0.0079], whereas comparing each epoch within females results in significance with Heat End over Before, Heat Start, and After [Before vs. Heat Start, D(136) = 0.2647, p value = 0.0135; Before vs. Heat End, D(136) = 0.4558, p value = 7.372e-07; Heat Start vs. Heat End, D(136) = 0.2352, p value = 0.0386; Heat Start vs. After, D(136) = 0.2941, p value = 0.0041; Heat End vs. After, D(136) = 0.3970, p value = 2.634e-05]). **(I)** Neuronal coactivity as measured by mean proportion of active cells (Two-way mixed-model ANOVA with Sidak’s post hoc: no Sex × Epoch interaction [F(3, 18) = 0.5414, p = 0.6601] or main effect of Sex [F(1, 6) = 0.9203, p = 0.3744]; main effect of Epoch [F(1.840, 11.04) = 4.163, p = 0.0475]). Data are shown as mean ±SEM. *p < 0.05; **p < 0.01; ***p < 0.001, ****p < 0.0001.

**Figure 2.**
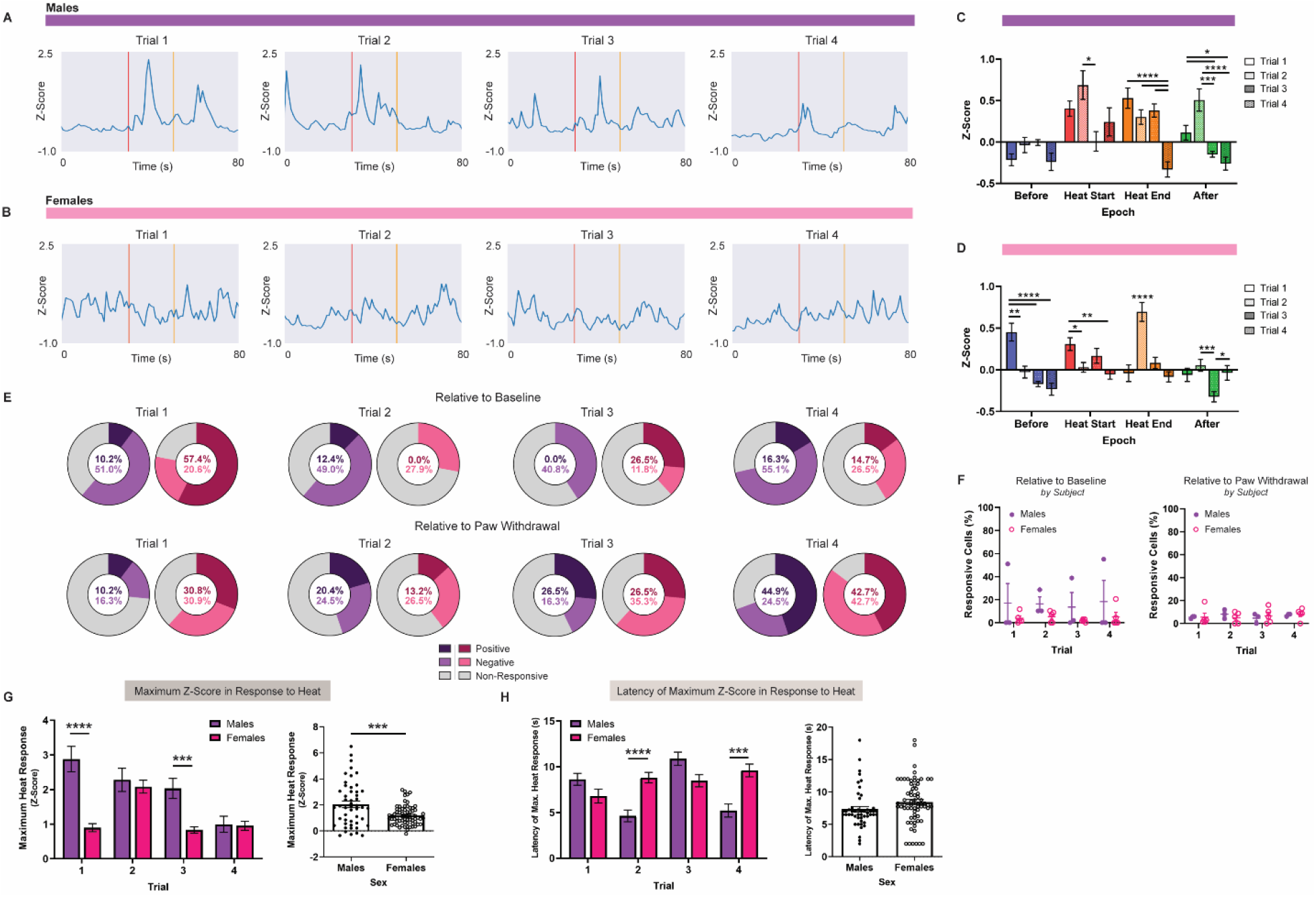
Progression of BNST^CRF+^ Activity Across Pain Exposure Trials. **(A-B)** Average BNST^CRF+^ traces of **(A)** male (n = 49 cells / 7-30 cells per mouse / 3 mice total) and **(B)** female (n = 68 cells / 8-19 cells per mouse / 5 mice total) subjects by trial. Within the representative time window, heat exposure starts at 30 seconds (first line [red]) with the maximum possible heat duration indicated at 50 seconds (second line [orange]). **(C-D)** Average z-score by epoch and trials for **(C)** male and **(D)** female subjects (<u>Males</u> = Two-way repeated measures [RM] ANOVA with Sidak’s post hoc: Epoch × Trial interaction [F(9, 576) = 7.373, p < 0.0001], main effect of Epoch [F(2.477, 475.5) = 21.42, p < 0.0001] and Trial [F(3, 192) = 8.695, p < 0.0001]; <u>Females</u> = Two-way RM ANOVA with Sidak’s post hoc: Epoch × Trial interaction [F(9, 804) = 17.04, p < 0.0001], main effect of Epoch [F(2.203, 590.4) = 13.95, p < 0.0001] and Trial [F(3, 268) = 6.913, p = 0.0002]). **(E)** Percentage of pain responsive cells for male (purple) and female (magenta) subjects were determined relative to epochs surrounding the onset of heat exposure and paw withdrawal across trials 1-4. Positive, negative, and non-responsive cells are indicated by darker purple/magenta, lighter purple/magenta, and grey respectively (Wilcoxon rank-sum, p < 0.05). **(F)** Proportion of pain responsive cells in individual subjects by trial and sex relative to the epochs surrounding (Left) heat onset (Two-way mixed-model ANOVA with Sidak’s post hoc: no Trial × Sex interaction [F(3, 18) = 0.0466, p = 0.9862] or main effect of Trial [F(1.092, 6.553) = 0.4473, p = 0.5435] and Sex [F(1, 6) = 1.510, p = 0.2651]) and (Right) paw withdrawal (Two-way mixed-model ANOVA with Sidak’s post hoc: no Trial × Sex interaction [F(3, 18) = 0.3670, p = 0.7777] or main effect of Trial [F(2.740, 16.44) = 0.3368, p = 0.7818] and Sex [F(1, 6) = 0.0008, p = 0.9776]). **(G)** (Left) Maximum z-score in response to heat by trial and sex (Two-way mixed-model ANOVA with Sidak’s post hoc: Trial × Sex interaction [F(3, 345) = 14.96, p < 0.0001], main effect of Trial [F(2.609, 300) = 20.28, p < 0.0001] and Sex [F(1, 115) = 13.69, p = 0.0003]). (Right) Average maximum z-score in response to heat by trial and sex (unpaired t-test: t(115) = 3.70, p = 0.0003). **(H)** (Left) Latency of maximum z-score in response to heat by trial and sex (Two-way mixed-model ANOVA with Sidak’s post hoc: Trial × Sex interaction [F(3, 345) = 16.90, p < 0.0001], main effect of Trial [F(2.717, 312.5) = 7.890, p < 0.0001], no main effect of Sex [F(1, 115) = 3.20, p = 0.0763]). (Right) Average latency of maximum z-score in response to heat stimulus by trial and sex (unpaired t-test: t(115) = 1.789, p = 0.0763). Data are shown as mean ±SEM. *p < 0.05; **p < 0.01; ***p < 0.001, ****p < 0.0001.

**Figure 3.**
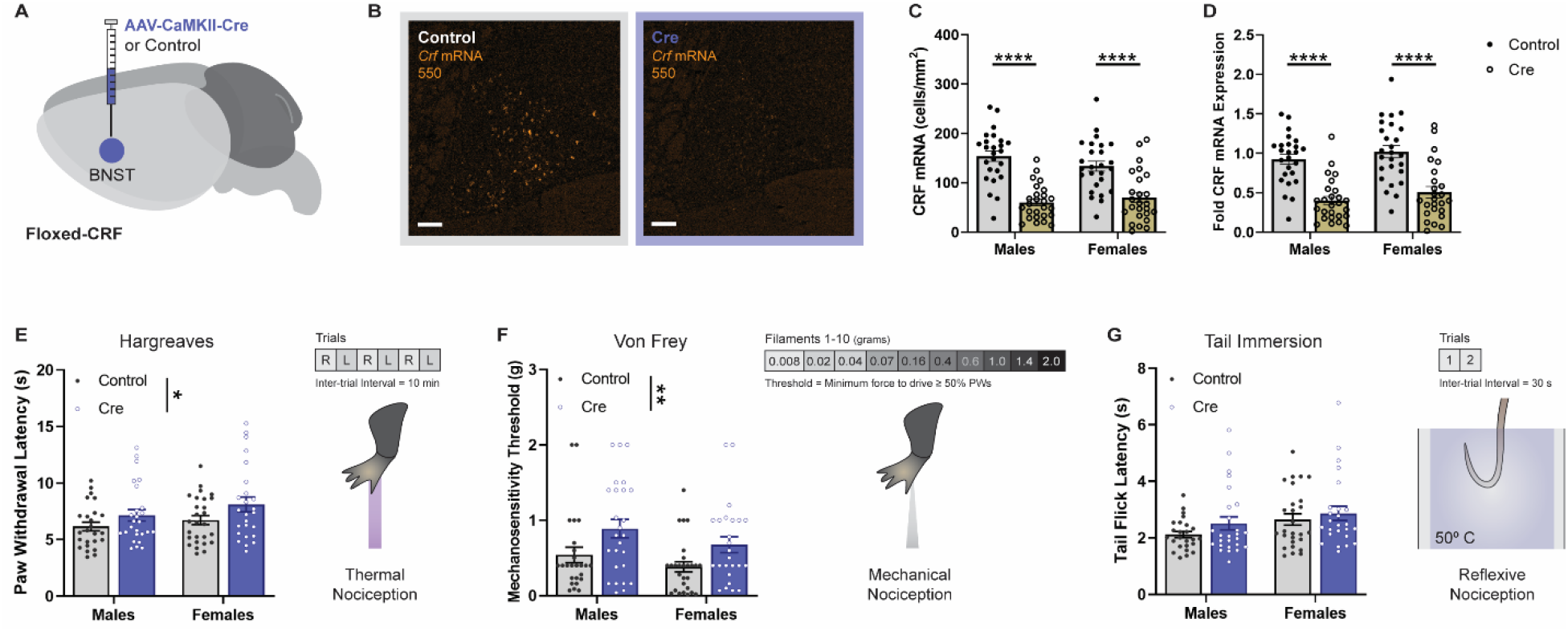
CRF Deletion in BNST Reduces Nociceptive Sensitivity. **(A)** Diagram of Cre-dependent approach for CRF deletion in BNST. **(B)** Representative histology of CRF mRNA expression in BNST of Floxed-CRF mice following virus infusion. **(C-D)** Quantification of CRF mRNA expression in BNST of male (n = 25-26) and female (n = 25-26) subjects by (C) cells/mm^2^ (Two-way ANOVA with Tukey’s post hoc: no Sex × Virus interaction [F(1, 98) = 2.548, p =0.1136] or main effect of Sex [F(1, 98) = 0.2667, p = 0.6067]; main effect of Virus [F(1, 98) = 66.74, p < 0.0001]) and (D) Fold change compared to control (Two-way ANOVA with Tukey’s post hoc: Two-way ANOVA: no Sex × Virus interaction [F(1, 98) = 0.007260, p =0.9323] or main effect of Sex [F(1, 98) = 2.239, p = 0.1378]; main effect of Virus [F(1, 98) = 60.23, p < 0.0001]). **(E-G)** Pain sensitivity of male (n = 25-26) and female (n = 25-27) subjects with schematic of behavioral test for **(E)** thermal (Two-way ANOVA with Tukey’s post hoc: no Sex × Virus interaction [F(1, 99) = 0.1585, p =0.6914] or main effect of Sex [F(1, 99) = 2.385, p = 0.1257]; main effect of Virus [F(1, 99) = 5.677, p = 0.0191]), **(F)** mechanical (Two-way ANOVA with Tukey’s post hoc: no Sex × Virus interaction [F(1, 99) = 0.07088, p =0.7906] or main effect of Sex [F(1, 99) = 3.297, p = 0.0724]; main effect of Virus [F(1, 99) = 10.05, p = 0.0020]), and **(G)** reflexive (Two-way ANOVA with Tukey’s post hoc: no Sex × Virus interaction [F(1, 99) = 0.1933, p =0.6611] or main effect of Virus [F(1, 99) = 2.224, p = 0.1391]; main effect of Sex [F(1, 99) = 4.631, p = 0.0338]) nociception. Data are shown as mean ±SEM. *p < 0.05; **p < 0.01; ***p < 0.001, ****p < 0.0001.

#### Von Frey

Subjects were held in Plexiglas boxes on a custom-made elevated metal wire surface (90 × 20 × 30 cm) and habituated to the behavioral apparatus for 30-60 minutes to ensure stable readouts of nociceptive sensitivity. Nylon monofilaments of forces ranging from 0.008 to 2 grams (g) were applied to the hind paw using the simplified up-down method (SUDO) described in Bonin *et al.* (2014). Starting with a mid-range force (0.16 g), the filament was applied to the mid-plantar surface of the hind paw for ten trials, then repeated with ascending or descending forces depending on the number of paw withdrawals. Withdrawal thresholds were defined as the minimum force filament that elicits a withdrawal reflex for ≥50% of the trials.

#### Tail Immersion

Following a 30-minute habituation to the behavioral room, subjects were restrained in Wypall fold wipers (Kimberly-Clark, Irving, TX) and tails were exposed to 50° C water in the test apparatus (Isotemp 110 Water Bath; Fisher Scientific, Hampton, NH). The tail flick latency was measured in two consecutive trials, where readings were taken 1 cm apart on the tail. The two latencies were then averaged together to indicate the reflexive nociceptive threshold of each subject. A cutoff time of 10 seconds was set to minimize tissue damage.

#### Hot Plate

Subjects were positioned in a custom-made cylindrical Plexiglas container on an IITC Hot Plate Analgesia Meter (IITC Life Science, Woodland Hills, CA). To elicit sensory-discriminative and affective-motivational behaviors associated with pain, subjects were exposed to a 55° C black anodized aluminum plate (11” × 10.5” × ¾”) for 45 seconds. This sustained version of the hot plate test extends the traditional cutoff time of 20 seconds to 45 seconds to improve readout for behaviors beyond latency to the first paw withdrawal. Behaviors like paw withdrawal (rapid flicking of the limb), paw attending (licking of the limb), paw guarding (intentional lifting of the limb for protection), and escape jumping (propelling into the air in an attempt to leave the apparatus) reflect the extent to which subjects will engage in reflexive responses versus motivated behaviors to reduce the aversive aspects of pain (Corder et al., 2017). Computer-integrated video cameras were used to record each session, allowing for blinded manual scoring of pain-related behaviors elicited by hot plate exposure. The behavioral apparatus was generously provided by the UNC Mouse Behavioral Phenotyping Laboratory.

#### Open Field

Subjects were given 30 minutes to habituate to the behavioral room before being placed into a white Plexiglas open field (50 × 50 × 25 cm), where they could freely explore the arena for 10 minutes. The center of the open field was defined as the central 25% of the arena, where light levels were approximately 30 lux. Tracking of subject location and activity was collected with EthoVision (Noldus Information Technologies, Wageningen, Netherlands).

#### Elevated Plus Maze

Following a 30-minute habituation period in the behavioral room, subjects were placed into the center of an elevated plus maze (EPM) and allowed to freely explore the arena for 10 minutes. During testing, subjects were able to explore two open arms (75 × 7 cm) and two closed arms (75 × 7 × 25 cm) that were bounded by a central area (7 × 7 × 25 cm). Light levels were set to approximately 15 lux to promote avoidance behaviors, which were determined by tracking subject location and activity with EthoVision (Noldus Information Technologies, Wageningen, Netherlands) and measuring time spent in the open and closed arms.

For all experiments, researchers were blinded to genotype/virus treatment conditions of each mouse.

### Ca^2+^ imaging with miniature microscope

#### Surgery

In CRF-Cre mice, a custom-prepared AAVDJ-EF1a-DIO-GCaMP6s virus (UNC Vector Core) was administered unilaterally in the dorsal region of BNST (250 nl in right hemisphere; relative to bregma: ML ±0.90 mm, AP +0.23 mm, DV −4.35 mm). GRIN lenses (0.6 mm diameter/7.3 mm length; Inscopix, Palo Alto, CA) were implanted approximately 200 μm over the virus injection site (relative to bregma: ML ±0.90 mm, AP 0.23 mm, DV −4.15 mm) and secured with a dental cement headcap. To promote a stable recovery following this extensive surgical procedure, subjects were given daily injections of meloxicam (5 mg/kg, s.c.) for four consecutive days. Approximately four weeks after the initial surgery and recovery, baseplates (V2; Inscopix, Palo Alto, CA) were placed above the GRIN lens to stabilize the attachment of the miniature microscope. The baseplate attachment was performed in a visually guided manner, where the miniature microscope was incrementally lowered towards the implanted GRIN lens to identify the optimal distance needed to achieve a field-of-view (FOV) with the maximum number of fluorescent cells. The baseplate was then incorporated into the existing headcap with additional dental cement. Finally, baseplate covers were secured to the top of the headcap in order to protect the miniature microscope attachment site from environmental contaminants that may obscure the imaging FOV.

#### Behavior

Subjects were tested in the Hargreaves assay 6-8 weeks after the initial surgical procedure and 2-4 weeks after the baseplate procedure. Prior to testing, each subject was anesthetized with isoflurane (1-3%) in oxygen (1-2 l/min) in order to attach the miniature microscope to the baseplate. After two consecutive days of habituation to the miniature microscope, subjects were given a 15-minute tethering session in their respective home cages and a 30-minute tethering session in the Hargreaves apparatus. At the end of the tethering session, a 5-minute imaging session for baseline calcium transients was recorded.

During testing, the mid-plantar surface of each hind paw was exposed to a series of noxious heat trials with 10-minute inter-trial intervals. Each subject was exposed to 4 trials of heat (I15 on the IITC Plantar Analgesia Meter [IITC Life Science, Woodland Hills, CA] respectively), with radiant heat exposures sequentially alternating between left (contralateral) and right (ipsilateral) paws. For each trial, calcium transients were recorded with a minute window prior to onset and following the offset of heat. Specific paw withdrawal latencies for each subject were used instead of a fixed duration to accurately capture individual nociceptive thresholds. Cutoff times of 20 seconds were set to prevent excessive tissue damage.

All recordings of behavior in the Hargreaves assay were performed with Media Recorder and a computer-integrated video camera (#PC212XP, Miniature CCD Camera Series). Simultaneous recordings of BNST^CRF+^ calcium transients were acquired using the nVista data acquisition software (Inscopix, Palo Alto, CA) at 20 fps (Exp Time: 50 ms, Gain: 4-7, Ex. LED Power: 1).

#### Preprocessing

Using the Inscopix Data Processing Software (IDPS; Inscopix, Palo Alto, CA), recordings were cropped, temporally downsampled to 5 fps, and motion corrected. ΔF/F was then acquired to normalize each pixel value in the recording to a baseline value (as determined by mean frame). To identify temporally and spatially unique ROIs, we used PCA/ICA to detect putative cell bodies in the imaging FOV. After manually removing ROIs that did not correspond to cells, traces and events were exported as CSV files. Preprocessed ΔF/F recordings were further transported into maximum intensity projection images and movie files for representative depictions of the data.

#### Data Analysis

Programmatic analysis was applied to extracted traces from IDPS using custom-written code in Python. Briefly, we transformed ΔF/F of baseline and test traces into z-scores for each subject using the formula: *Z* = × - *μ* / *δ*, where the number of standard deviations (*δ*) that raw values (*x*) diverge from baseline values (*μ*) is determined. We then separated traces by epoch (*Before*, *Heat Start*, *Heat End*, *After*) for each subject and trail to determine averages by sex. Epoch timing was determined by dividing the duration of heat exposure into halves, so that each time block represented an equal amount of time for specific trials and subjects. Analysis was then performed for various combinations of data dimensionality based on these times.

We specifically compared the activity of individual male and female BNST^CRF+^ neurons for the following metrics:

–*Average Z-Score Across Trials by Epoch*: Mean z-score of epochs averaged across trials.
–*Average Z-Score by Trial and Epoch*: Mean z-score of epochs for each trial.
–Average *Neuronal Coactivity by Epoch*: Mean proportion of synchronized cell responses for each epoch. Fractions of responsive cells were calculated at every time frame for individual subjects. The average fraction of responses was then sorted by epoch for comparison.
–*Cumulative Distribution Function Across Trials by Epoch*: Determine the distribution of z-scores averaged across trials by binning the frequency of activity within each epoch.
–*Percent Responsiveness Across Trials*: Determine the proportion of cells that exhibit a statistically significant change in average z-scores during heat exposure by using a Wilcoxon rank-sum test for each neuron. Two notable events define comparisons of heat responsive neurons with the activity in surrounding epochs: “Heat Onset” (i.e. *Heat On* [*Heat Start* + *Heat End*] vs. *Before*) and “Paw Withdrawal” (i.e. *Heat On* vs. *After*). Positive differences in z-score correspond to activation in response to noxious heat exposure relative to surrounding epochs, while negative differences indicate a relative reduction in activity. Percentage of responsive cells was calculated by factoring in every neuron for each trial. Separate analyses restricted proportions of responsive cells by individual subjects.
–*Percent Responsiveness by Trials*: Determine the change between average z-scores of *Heat On* (i.e. *Heat Start* and *Heat End*) and surrounding epochs for each trial, with positive differences corresponding to activation in response to noxious heat exposure, and negative differences corresponding to inactivation. Percentage of responsive cells was calculated by taking the proportion of neurons exhibiting statistically different *Heat On* activity compared to *Before* and *After* in individual subjects and averaging these values by subject within each trial.
–*Spike Magnitude by Trials*: Identify size of maximum value in *Heat On* (i.e. *Heat Start* and *Heat End*) epoch for each trial.
–*Spike Latency by Trials*: Identify time stamps for maximum value in *Heat On* (i.e. *Heat Start* and *Heat End*) epoch for each trial.

More information on the Python scripts used for analysis can be requested by contacting.

### In situ hybridization

Following isoflurane anesthetization and rapid decapitation, the brains of Floxed-CRF mice were collected and placed on aluminum foil, where they were immediately frozen on dry ice and stored in a −80° C freezer. Using a Leica CM3050 S cryostat (Leica Microsystems, Wetzlar, Germany), coronal sections of BNST (12 μm thickness) were obtained and directly mounted onto Superfrost Plus slides (Fisher Scientific, Hampton, NH), then kept at −80° C. In order to fluorescently label *Crf* mRNA in the BNST, slices were preprocessed with 4% PFA and protease reagent, incubated with target probes for mouse *Crf* (#316091, RNAscope Probe – Mm-Crh; Advanced Cell Diagnostics, Newark, CA), then fluorescently labeled with a probe targeting the corresponding channel for the peptide (Crf in 550; Advanced Cell Diagnostics, Newark, CA).

The processed slides were then covered using Vecta-Shield Mounting Medium with DAPI in preparation for imaging.

### Confocal Microscopy

All fluorescent images were acquired with the Zeiss 800 Upright confocal microscope and ZenBlue software (Carl Zeiss AG, Oberkochen, Germany), with equipment access granted through the Hooker Imaging Core at UNC Chapel Hill. Validation of virus expression/injection site, GRIN lens placement, and immunoreactivity were accomplished with tiled and serial z-stack images obtained through a 20× objective (3 × 3 tile, 2 μm optical slice thickness, 12 μm total thickness). Images were processed in FIJI (Schindelin et al., 2012) for manual counting and ZenBlue (Carl Zeiss AG, Oberkochen, Germany) for area measurements in each ROI.

### Statistical analysis

Single-variable comparisons were made using unpaired t-tests. Group comparisons were made using two-way ANOVA, two-way repeated measures ANOVA, or two-way mixed-model ANOVA depending on the number of independent and within-subjects variables in a data set. Following significant interactions or main effects, post-hoc pairwise t-tests were performed and corrected using Sidak’s or Tukey’s post-hoc tests to control for multiple comparisons. Results of statistical testing are reported in figure legends with significance indicated through markers on figures. Data are expressed as mean ± standard error of the mean (SEM), with significance for *p* values below 0.05 (* p < 0.05, ** p < 0.01, *** p < 0.001, **** p < 0.0001). All data were analyzed and visualized with standard statistical software packages from GraphPad Prism 8 (GraphPad Software, San Diego, CA) and SciPy (Virtanen et al., 2020).

Details on the statistical tests used:

Unpaired t-test used for *Figures 1B, 2G-H*.

Wilcoxon rank-sum test used for *Figures 1D, 2E*.

Kolmogorov–Smirnov test used for *Figure 1H*.

Area under the curve used for *Figures 4D-E*.

Two-way ANOVA with Tukey’s post hoc used for *Figures 1E, 3C-G, 4B-C, 5A-D*.

Two-way mixed-model ANOVA with Tukey’s post hoc used for *Figures 1G*.

Two-way mixed-model ANOVA with Sidak’s post hoc used for *Figures 1B, 1H, 2F-H*.

Two-way repeated measures ANOVA with Sidak’s post hoc used for *Figures 2C-D*.

**Figure 4.**
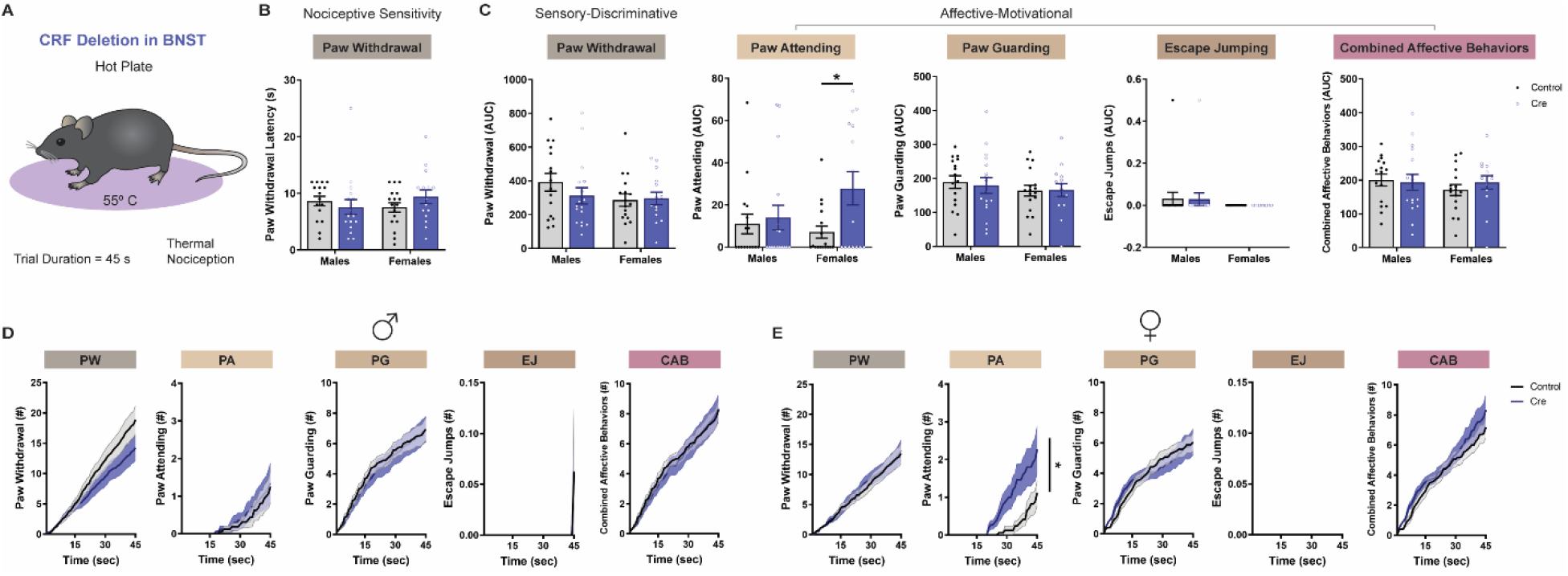
CRF Deletion in BNST Differentially Alters Sensory-Discriminative and Affective-Motivational Behaviors. **(A)** Schematic of hot plate test. **(B)** Thermal nociceptive sensitivity of male (n = 16-17) and female (n = 15-17) Floxed-CRF mice, as defined by latency to first paw withdrawal for each mouse (Two-way ANOVA with Tukey’s post hoc: no Sex × Virus interaction [F(1, 61) = 1.853, p =0.1785] or main effect of Sex [F(1, 61) = 0.1124, p = 0.7386] and Virus [F(1, 61) = 0.1524, p = 0.6976]). **(C)** Quantification of sensory-discriminative and affective-motivational behaviors throughout the 45-second exposure to hot plate. Averaged AUC of paw withdrawal (Two-way ANOVA with Tukey’s post hoc: no Sex × Virus interaction [F(1, 61) = 1.072, p =0.3047] or main effect of Sex [F(1, 61) = 1.885, p = 0.1747] and Virus [F(1, 61) = 0.6208, p = 0.4338]), paw attending (Two-way ANOVA with Tukey’s post hoc: no Sex × Virus interaction [F(1, 61) = 2.605, p =0.1117] or main effect of Sex [F(1, 61) = 0.8298, p = 0.3659]; main effect of Virus [F(1, 61) = 4.695, p = 0.0342]), paw guarding (Two-way ANOVA with Tukey’s post hoc: no Sex × Virus interaction [F(1, 61) = 0.09333, p =0.7610] or main effect of Sex [F(1, 61) = 1.026, p = 0.3152] and Virus [F(1, 61) = 0.04448, p = 0.8337]), escape jumping (Two-way ANOVA with Tukey’s post hoc: no Sex × Virus interaction [F(1, 61) = 0.001778, p =0.9665] or main effect of Sex [F(1, 61) = 1.936, p = 0.1691] and Virus [F(1, 61) = 0.001778, p = 0.9665]), and combined affective behaviors (Two-way ANOVA with Tukey’s post hoc: no Sex × Virus interaction [F(1, 61) = 0.5696, p =0.4533] or main effect of Sex [F(1, 61) = 0.5631, p = 0.4559] and Virus [F(1, 61) = 0.1580, p = 0.6924]) are shown, with corresponding cumulative distribution functions displayed in **(D-E)**. **(D-E)** Sensory-discriminative and affective-motivational behaviors are exhibited across time with cumulative distribution functions for **(D)** male and **(E)** female subjects (paw withdrawal: CON (M): 391.9 [341.0-442.9], CRE (M): 311.7 [263.2-360.2], CON (F): 285.9 [249.1-322.7], CRE (F): 296.8 [261.1-332.5]; paw attending: CON (M): 11.00 [4.672-17.33], CRE (M): 14.03 [6.087-21.97], CON (F): 7.147 [2.442-11.85], CRE (F): 27.87 [17.95-37.78]; paw guarding: CON (M): 189.3 [171.7-207.0], CRE (M): 179.3 [156.7-201.9], CON (F): 163.8 [147.6-180.0], CRE (F): 165.6 [147.8-183.4]; escape jumping: CON (M): 0.03125 [0-0.2762], CRE (M): 0.2941 [0-0.2671], CON (F): 0 [0-0], CRE (F): 0 [0-0]; combined affective behaviors: CON (M): 200.4 [183.4-217.3], CRE (M): 193.4 [170.1-216.7], CON (F): 170.9 [154.3-187.5], CRE (F): 193.5 [173.8-213.1]). Data are shown as mean ±SEM. *p < 0.05; **p < 0.01; ***p < 0.001, ****p < 0.0001.

## Results

### BNST^CRF+^ neurons are endogenously recruited by noxious heat

To assess how BNST^CRF+^ neurons are modulated by pain, we used a head-mounted miniature microscope to track *in vivo* somatic calcium activity of individual *Crf+* neurons in the BNST following hind paw exposure to noxious heat (Figure 1). By implanting a GRIN lens and injecting an adeno-associated virus carrying Cre-inducible GCaMP6s (AAVDJ-EF1a-DIO-GCaMP6s) in the right hemisphere BNST of adult male and female CRF-Cre mice (Figure 1A), we were able to continuously monitor BNSTCRF+ activity during repeated trials of noxious heat exposure. Since the length of exposure varied by trial, we analyzed activity across epochs relative to heat exposure: *Before*, *Heat Start*, *Heat End*, and *After* (Figures 1C and 1E). During the simultaneous measurement of paw withdrawal latencies in male (n = 3) and female (n = 5) mice (Figure 1B), comparisons of BNST^CRF+^ activity averaged across four trials of heat exposure revealed sex-specific recruitment by epoch, with higher fold calcium changes observed for *Heat Start* and *Heat End* in males (n = 49 cells; 7-30 cells per mouse) and *Heat End* in females (n = 68 cells; 8-19 cells per mouse) (Figures 1G–1H;main effect of Epoch: F_2.153, 247.7_ = 26.31, p < 0.0001). Average z-scores of *Before* and *Heat Start* epochs differed by sex, with males showing a larger magnitude of BNST^CRF+^ activity in the earlier phases of stimulus exposure than females (Figure 1G;main effect of Sex: F_1, 115_ = 13.20, p = 0.0004). Coactivation of neuronal activity, as defined by the average fraction of responsive cells for each frame of an epoch, also differed in the earlier phases of stimulus exposure, with females exhibiting increased BNST^CRF+^ synchrony between the transition from *Before* to *Heat Start* (Figure 1I;main effect of Epoch: F_1.840, 11.04_ = 4.163, p = 0.0475). Notably, while there were sex differences in the magnitude and synchrony of responses when examining average z-scores, the proportion of cells exhibiting time-locked changes to heat exposure did not differ between male and female subjects (Figure 1C). These results indicate that BNSTCRF+ neurons are recruited during thermal nociception in a manner where the magnitude of activation appears greater in males, while synchronous activation is more prominent in females.

We next evaluated BNST^CRF+^ activity on a trial-by-trial basis to determine how neuronal responses to noxious heat are impacted by repeated exposure **(Figures 2A–2B)**. Higher z-scores were generally observed in the first two trials of *Heat Start* and *Heat End* epochs **(Figures 2C–2D)**. Although male mice exhibited averaged BNST^CRF+^ responses with a greater magnitude of activation than females **(Figures 2G**; t_115_ = 3.70, p = 0.0003), maximum heat responses were reduced throughout the progression of trials for males **(Figure 2G**; main effect of Sex: F_1, 115_ = 13.69, p = 0.0003). By contrast, the latency of maximum response became more delayed with additional trials for female mice **(Figure 2H**; Trial X Sex interaction: F_3, 345_ = 16.90, p < 0.0001), resulting in a trend for later peak times following heat exposure **(Figure 2H**; t_115_ = 1.789, p = 0.0763). Tracking the percentage of heat responsive BNST^CRF+^ neurons across trials, we observed a combination of positive and negative changes in activity that varied in distribution with repeated exposures **(Figure 2E)**. However, when restricted to cell proportions by subject, the percentage of heat responsive BNST^CRF+^ neurons for each trial did not significantly differ by trial progression or sex **(Figure 2F)**. These data suggest that multiple exposures to acute thermal nociception can drive distinct changes in the magnitude and timing of cell responses, but not the total proportion of responsive cells among individual subjects, for male and female mice.

Together, our *in vivo* imaging results indicate that BNST^CRF+^ neurons are activated by noxious heat in male and female mice.

### Genetic deletion of CRF from BNST reduces pain sensitivity

Given the heterogenous nature of peptide expression in the BNST and a fundamental lack of understanding as to what conditions drive CRF release, it was unclear whether the observed BNST^CRF+^ responses to noxious heat in our imaging experiments would translate to a functional role for CRF in the modulation of pain-related behaviors. To address this issue, we genetically deleted CRF in the BNST by bilaterally injecting an adeno-associated virus carrying Cre (AAV5-CaMKIIa-Cre-GFP) or a control virus (AAV5-CaMKIIa-GFP) in the BNST of adult male and female Floxed-CRF mice **(Figure 3A)**. This approach resulted in an approximate 50-60% reduction in *Crf* mRNA-positive neurons in the BNST **(Figures 3B–3D**; Crf mRNA: main effect of Virus: F_1, 98_ = 66.74, p < 0.0001; Fold Crf mRNA: main effect of Virus: F_1, 98_ = 60.23, p < 0.0001). Using the Hargreaves and Von Frey tests, we found that CRF deletion reduced nociceptive sensitivity compared to control mice **(Figures 3E–3F**; Hargreaves: main effect of Virus: F_1, 99_ = 5.677, p = 0.0191; Von Frey: main effect of Virus: F_1, 99_ = 10.05, p = 0.0020). Notably, the extent of reduction for *Crf* mRNA expression and pain sensitivity did not differ between male and female subjects **(Figures 3C–3F)**. By contrast, reflexive responses in the tail immersion test revealed lower nociceptive thresholds in males, but no change in sensitivity following CRF deletion **(Figure 3G**; Tail Immersion: main effect of Sex: F_1, 99_ = 4.631, p = 0.0338). These results show that a deficiency of CRF in the BNST increases supraspinal pain thresholds, suggesting that the presence of CRF may be necessary to preserve the sensory components of pain for both male and female mice.

### Genetic deletion of CRF from BNST differentially impacts the sensory and affective-motivational components of pain

Considering that CRF signaling in the BNST plays an important role in multiple components of the pain experience, we tested the same CRF deletion and control mice in an extended hot plate test (Corder et al., 2017). Differing from a standard hot plate protocol that terminates after the first paw withdrawal or a relatively brief predetermined cut-off time, this prolonged form of the test lasts 45 seconds to promote the expression of both sensory-discriminative (e.g. paw withdrawal [rapid flicking of the limb]) and affective-motivational (e.g. paw attending [licking of the limb], paw guarding [intentional lifting of the limb for protection], and escape jumping [propelling into the air in an attempt to leave the apparatus]) behaviors **(Figure 4A).** Although genetic deletion of CRF in the BNST did not alter sensory-discriminative responses to the hot plate, as measured by paw withdrawal latency **(Figure 4B)** and cumulative paw withdrawal behaviors **(Figure 4C)**, or affective-motivational responses such as paw guarding and escape jumping **(Figure 4C)**, there was a marked increase in paw attending for Cre-treated female mice **(Figures 4C–4E**; main effect of Virus: F_1, 61_ = 4.695, p = 0.0342). This sex-specific change in paw attending suggests a divergent role for CRF in the BNST and the sensory-discriminative/affective-motivational components of a prolonged, inescapable thermal nociceptive stimulus. Remarkably, reducing CRF expression in the BNST did not impact other pain-independent affective-motivational behaviors such as avoidance **(Figures 5A and 5C)** and locomotor **(Figures 5B and 5D)** behaviors in the open field **(Figures 5A–5B)** and elevated plus maze **(Figures 5C–5D)**, indicating that the aversive contributions of CRF in the BNST are specific to pain-related contexts like the hot plate.

**Figure 5.**
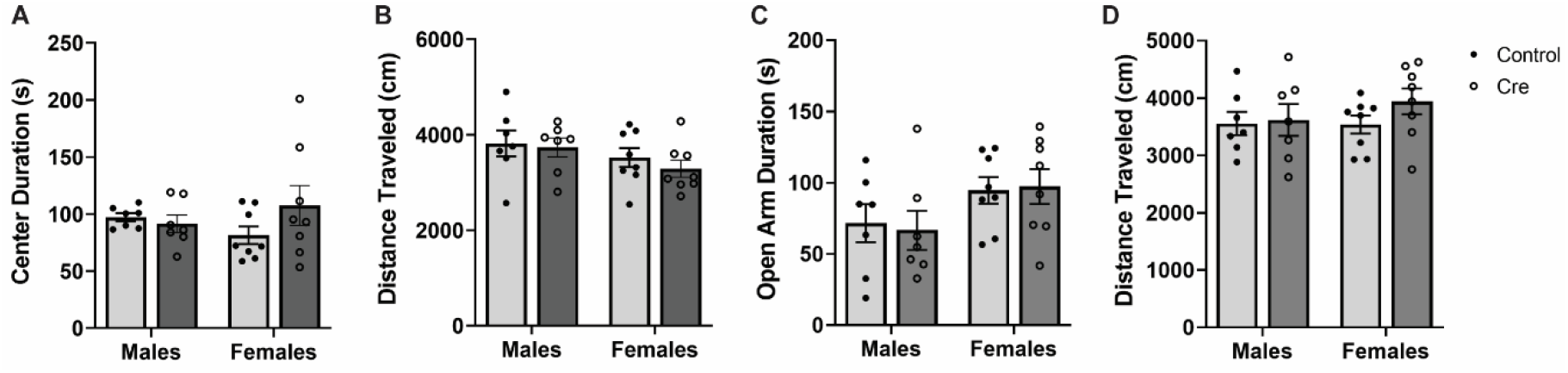
CRF Deletion in BNST Does Not Alter Avoidance Behaviors. **(A)** Avoidance behaviors in male (n = 7) and female (n = 8) Floxed-CRF mice, as measured by duration in the center (seconds) of the open field (Two-way ANOVA with Tukey’s post hoc: no Sex × Virus interaction [F(1, 26) = 2.098, p = 0.1594], no main effect of Sex [F(1, 26) = 0.0001, p = 0.9911] and Virus [F(1, 26) = 0.8511, p = 0.3647]). **(B)** Locomotor behaviors in male (n = 7) and female (n = 8) subjects, as measured by distance traveled (cm) in the open field (Two-way ANOVA with Tukey’s post hoc: no Sex × Virus interaction [F(1, 26) = 0.1251, p = 0.7265], no main effect of Sex [F(1, 26) = 3.015, p = 0.0943] and Virus [F(1, 26) = 0.5665, p = 0.4584]). **(C)** Avoidance behaviors in male (n = 7) and female (n = 8) subjects, as measured by duration in the open arms (seconds) of the elevated plus maze (Two-way ANOVA with Tukey’s post hoc: no Sex × Virus interaction [F(1, 26) = 0.0995, p = 0.7549], main effect of Sex [F(1, 26) = 4.908, p = 0.0357], no main effect of Virus [F(1, 26) = 0.0083, p = 0.9280]). **(D)** Locomotor behaviors in male (n = 7) and female (n = 8) subjects, as measured by distance traveled (cm) in the elevated plus maze (Two-way ANOVA with Tukey’s post hoc: no Sex × Virus interaction [F(1, 26) = 0.6221, p = 0.4374], no main effect of Sex [F(1, 26) = 0.4944, p = 0.4882] and Virus [F(1, 26) = 1.161, p = 0.2911]). Data are shown as mean ±SEM. *p < 0.05; **p < 0.01; ***p < 0.001, ****p < 0.0001.

## Discussion

Pain is a multi-faceted experience that promotes aversion and necessitates adaptive coping strategies to mitigate harm. In the present study, we identified a subpopulation of BNST neurons that can respond to and modulate pain-related behaviors through local expression of CRF. Utilizing genetically targeted manipulations of CRF-containing neurons and *Crf* itself in the BNST, we were able to demonstrate the dynamic recruitment of BNST^CRF+^ activity by noxious heat, as well as the modulatory role of CRF on pain in male and female mice. Our results support and expand upon earlier work on intra-BNST CRF signaling and pain (Tran et al., 2014; Minami, 2019), which has historically been limited to the role of CRF receptors in the BNST of male rodents, by uncovering previously unknown sex-specific contributions of local CRF production. These findings enhance our collective knowledge on nociceptive microcircuits in the BNST.

### BNST^CRF+^ neurons dynamically respond to noxious heat across trials in a sex-specific manner

Our imaging experiments determined that BNST^CRF+^ neurons generate greater somatic Ca^2+^ activity in the presence of noxious heat. Since the greatest increases in activity were observed at the *Heat Start* and *Heat End* epochs for males and the *Heat End* epoch for females, we conclude that BNST^CRF+^ neurons are activated by noxious heat in both sexes. Sex differences at the *Before* and *Heat Start* epochs, where the initial detection of the stimulus elicited greater neuronal responses in males, additionally suggest that neuronal representations of pain differ by sex in the early phases of heat exposure. This difference in early phase processing may indicate a higher probability of detecting the nociceptive stimulus, since males exhibited trends for BNST^CRF+^ neurons with faster maximum heat response times and greater magnitudes of activation. By contrast, females exhibited greater BNST^CRF+^ coactivity changes in the early phases of heat exposure, with the most prominent increases in synchrony happening between the transition from *Before* to *Heat Start*. We posit that these disparities in early phase activity reflect sex-specific mechanisms of nociceptive processing. Considering that salience is an important predictor of pain sensitivity and analgesic efficacy (Borsook et al., 2013), it is possible that alterations in the timing and synchrony of BNST^CRF+^ responses will change nociceptive detection and contribute to pain sensitivity in a sex-specific manner. This is an intriguing possibility given that BNST neurons expressing *Crf* or the prepronociceptin gene have been reported to exhibit *in vivo* responses to motivationally salient and aversive stimuli such as predator odor (Giardino et al., 2018; Rodriguez-Romaguera et al., 2020). These observations support prior conclusions that the BNST encodes for salient threats that may cause physical or emotional harm (Somerville et al., 2010; Borsook et al., 2013; Grupe et al., 2013; Avery et al., 2016; Hermann et al., 2016; Minami, 2019).

Since salience can either enhance or diminish the importance of a nociceptive signal with experience (Borsook et al., 2013), we additionally evaluated whether BNST^CRF+^ responses to heat change with repeated exposure. Comparisons by sex revealed increasingly lower magnitudes of activity for males and slower maximum heat response times for females as trials advanced. These progressive changes in magnitude and latency suggest that BNST^CRF+^ neurons encode for novelty, a distinctive feature that contributes to the nociceptive impact of each stimulus exposure, where the extent of response decreases and becomes more delayed as subjects habituate to the experience. Clinical reports similarly show that repeated exposure to a nociceptive stimulus leads to dissociations in stimulus representation by the brain, with the BNST being implicated in the gating processes that determine the significance of environmental stimuli (Becerra et al.,1999; Herrmann et al., 2016). Reductions in BNST^CRF+^ activity may thus reflect the diminished salience of pain as stimulus exposures accumulate. In reference to related structures like the basolateral amygdala and the central nucleus of the amygdala [CeA], which exhibit neuronal responses that positively scale with increasing pain intensity and repeated nociception (Ji et al., 2010; Grewe et al., 2017; Yu et al., 2017; Corder & Ahanonu et al., 2019; for a non-responsive example, see Hua et al., 2020), BNST^CRF+^ responses are distinctly desensitized with repeated pain exposures. Interestingly, sex differences in the size and timing of responses did not translate to test performance, as thermal nociceptive sensitivity was comparable between male and female mice across trials. This suggests that the observed discrepancies in BNST^CRF+^ encoding between males and females are indicative of a sex-specific means to process pain rather than a mechanism to drive behavioral variation. Quantitative comparisons have shown that female subjects are more sensitive than males in 85.4% of rodent studies (Mogil et al., 2020), so our recordings of *in vivo* BNST^CRF+^ responses to various components of the pain experience offer unique insight on how comparable nociceptive sensitivity can be processed in a sex-specific manner.

Notably, we observed variability in BNST^CRF+^ responding on a single cell level for both sexes, where responses to heat were comprised of both increasing and decreasing z-scores compared to activity in the *Before* and *After* epochs. Previous data from our group has suggested that multiple populations of CRF neurons in the BNST differentially contribute to the regulation of affective behaviors (Marcinkiewcz and Mazzone et al., 2016), with BNST^CRF+^ neurons that form local inhibitory synapses playing a greater role in aversive states. Assuming that the magnitude of response is a reliable metric of activation, these data suggest a model where heightened BNST^CRF+^ activation is accompanied by increased inhibitory tone. Suppression of local GABAergic projection neurons has been shown to inhibit dopamine neurons in the ventral tegmental area (Marcinkiewcz and Mazzone et al., 2016, Takahashi et al., 2019), so it is possible that the rapid decreases in BNST^CRF+^ activity observed prior to paw withdrawal are providing a motivational signal to move away from the noxious stimulus. Although BNST^CRF+^ neurons exhibit local inhibitory connections, it should be noted that they also project to other regions involved in motivated behavior, such as the lateral hypothalamus, where activation of BNST^CRF+^ projections can induce conditioned place aversion (Dabrowska et al., 2016; Giardino et al., 2018). In future experiments, researchers should more rigorously examine the functional contributions of these varied BNST^CRF+^ responses to pain-related behaviors, with special consideration for how the proportion of pain responsive neurons and magnitude/timing of activity interact to inform function.

### CRF in the BNST modulates the expression of sensory and affective-motivational components of pain

While previous studies have found that pharmacological manipulation of CRF signaling in the BNST can modulate multiple aspects of pain-related behavior (Ide et al., 2013; Tran et al., 2012; 2014; Kaneko et al., 2016), the source of this CRF is unclear, as the BNST contains locally produced CRF and receives CRF input from the CeA (Sakanaka et al., 1986; Dong et al., 2001). In examining how local production of CRF can impact these pain-related behaviors, our work has revealed that both thermal and mechanical nociceptive sensitivity are impacted by CRF deletion in the BNST. These results are reminiscent of previously published effects showing that CRF_2R_ antagonism reduces mechanical and colonic nociception in stress-free conditions (Tran et al., 2014). However, similar pharmacological manipulations failed to alter nociceptive behaviors associated with formalin and only blocked conditioned place aversion (Ide et al., 2013; Kaneko et al., 2016). In the context of our results, these studies suggest that CRF signaling in the BNST selectively contributes to the sensory and affective-motivational components of pain based on the specific nociceptive stimulus and the context of its presentation. As others have hypothesized, the duration of a nociceptive stimulus and the capacity of an animal to prevent further harm by the stimulus (i.e. escapability) may be important factors in pain processing (Ryan et al., 1985; Bandler et al., 2000; Gandhi et al., 2017). If we categorize these known effects of CRF signaling in the BNST by phasic-acting/escapable nociceptive exposures like the Hargreaves and Von Frey tests versus tonic-acting/inescapable nociceptive exposures like formalin, we can posit that CRF is more likely to alter the sensory components of pain when challenged with shorter-lasting stimuli, while the affective-motivational components are more likely to be influenced by longer-lasting stimuli.

Contrary to the anti-nociceptive effects observed with acute nociceptive exposures in the Hargreaves and Von Frey tests, we found that CRF deletion in the BNST did not impact the sensory components of pain in the hot plate when measured by initial latency and cumulative number of paw withdrawals. Of the measures for affective-motivational behaviors, only paw attending was altered by CRF deletion, as females exhibited more licks to the site of injury in Cre-treated mice than controls. Unlike the Hargreaves and Von Frey tests, where exposure to the nociceptive stimulus terminates after a paw withdrawal or a fixed number of rapid prodding, the extended hot plate exposes subjects to heat for a duration where the thermal nociceptive stimulus is tonic-acting/inescapable compared to the standard version of the test. This change in the context of nociceptive stimulus presentation may explain why CRF deletion can generate seemingly contradictory phenotypes like sex-independent pain relief and sex-dependent exacerbation of an adaptive coping behavior. These divergent results possibly indicate a functional distinction in how CRF signaling in the BNST interacts with specific nociceptive stimuli as phasic- or tonic-acting stressors.

Interactions between nociceptive stimuli, the context of their presentation, and sex-specific adaptative behaviors may be explained by variable approaches to processing the salient features of an environment, as the same nociceptive experience may require differential engagement of CRF signaling in the BNST for male and female subjects. The BNST has been implicated in pain and salience through its role in hypervigilant threat monitoring, a form of sustained attention reserved for potentially harmful stimuli (Herrmann et al., 2016; Brinkmann et al., 2017; Jenks et al., 2020). Systemic administration of CRF produces impairments in sustained attention, with the largest reductions in responses to salient stimuli observed in the diestrus phase of the estrous cycle for female rats (Cole et al., 2016). Estrous cycle influence on sustained attention may thus translate to discrete processing of tonic-acting/inescapable nociceptive stimuli between males and females, where CRF deletion in the BNST enhances the detection of longer-lasting pain for females to increase active coping behaviors, while the same CRF manipulation is less susceptible to estrous influence when confronted with phasic-acting/escapable nociceptive stimuli, since the detection of acute pain may depend less on sustained attention (Sommerville et al., 2010; Borsook et al., 2013; Hermann et al., 2016). While the present study did not track fluctuations in the estrous cycle, it is an intriguing possibility for sex hormones to conditionally interact with CRF signaling based on salience, so that tonic-acting/inescapable contexts increase the likelihood of sex-specific responses to pain. This theoretical framework supports earlier work establishing sex differences in peptide distribution, receptor expression, and stress response as broad features of CRF signaling throughout the brain (Funabashi et al., 2004; Bangasser & Wiersielis, 2018; Uchida et al., 2019), with many of these phenotypes being the product of hormonal surges during development (Bangasser & Wiersielis, 2018). Whether similar sex-specific functions of CRF signaling apply to the BNST was not addressed in previous evaluations of CRF signaling and pain, as these studies were performed exclusively in male subjects (Deyama et al., 2007; Tran et al., 2012; 2014; Ide et al., 2013; Kaneko et al., 2016). By consequence, the findings described in the present study are the first to compare how local production of CRF in the BNST contributes to different modalities of pain in both sexes. Additional studies on the impact of the estrous cycle on nociceptive stressors of varying duration/escapability may be necessary to more fully understand how CRF signaling in the BNST contributes to discrete pain processing and modulation in male and female mice.

Although there were female-specific changes in active coping responses to pain, no differences in avoidance or locomotors behaviors were observed with CRF deletion in the BNST. Previous studies have shown that CRF infusion in the BNST is anxiogenic and produces conditioned place aversion, while pharmacological inhibition of CRF_1R_ but not CRF_2R_ in the BNST is anxiolytic (Sahuque et al., 2006). Furthermore, work from our lab has shown that chemogenetic inhibition of BNST^CRF+^ neurons reduces avoidance behavior in the open field (Pleil et al., 2015).

Our results demonstrate that genetic targeting of CRF production does not reduce avoidance behaviors, similar to the null effects observed in a previous study with CRF overexpression in the BNST (Sink et al., 2013), suggesting that there may be other factors in BNST^CRF+^ neurons driving these changes. Although there has yet to be a comprehensive investigation on BNST^CRF+^ co-expression and its relative contributions to avoidance behaviors, distinct anxiogenic roles have been identified for GABA, CRF, and other modulatory factors in the CeA (Pomrenze et al., 2019). It remains possible, however, that CRF plays a prominent role in avoidance behaviors, and that a greater reduction in *Crf* mRNA beyond 50%-60% is necessary to produce anxiolytic effects. Alternatively, it is feasible that anxiety-like states are more robustly regulated by external CRF contributions than local contributions within the BNST.

Taken together, our work raises important questions on the relationship between the dynamic activity of BNST^CRF+^ neurons and CRF release. Unlike classical neurotransmitters such as GABA, the kinetics of release and activity of neuropeptides can be much longer. Elegant work from the Minami group has demonstrated that CRF can be released in the BNST over the course of an hour following formalin injection (Ide et al., 2013), so it is possible that some of the changes we observed over time in our *in vivo* imaging studies were due to local release of CRF. This is a particularly interesting possibility in the context of chronic pain, since long-term injury has been reported to drive functional upregulations of CRF signaling and putative CRF activity in the BNST (Rouwette et al., 2012; Ide et al., 2013; Takahashi et al., 2019; Hara et al., 2020).

Interestingly, treating adult male human subjects with CRF has been shown to relieve pain during persistent inflammatory states such as post-operative pain but lack analgesic properties for healthy individuals (Hargreaves et al., 1987; Lautenbacher et al., 1999), suggesting that lasting pain conditions can change how CRF modulates pain. Future studies examining the relationship between chronic pain and *in vivo* dynamics of BNST^CRF+^ activity in response to noxious stimuli will therefore be essential for a comprehensive understanding of peptide function in pain processing and modulation.

To summarize, our findings have established a previously uncharacterized sex-specific role for CRF in the BNST and pain-related behaviors that will provide important insight into the aversive microcircuitry of the extended amygdala. We hope that this knowledge will present a viable path towards more equitable approaches to recognizing and managing pain for both sexes.

## Acknowledgements

This work was supported by grants from the National Institute of Health (NIH) and the National Institute on Alcohol Abuse and Alcoholism (NIAAA): T32GM007040 [W.Y.], F31AA027129 [W.Y.], and R21AA027460 [T.K.]. We would like to thank Mark Miller, Shay Neufeld, and Inscopix Data Services for their help with neuronal coactivity analysis, as well as Juan Song, Andrea G. Nackley, Kathryn J. Reissner, and Zoe A. McElligott for their support and critical feedback over the course of the project.

## Author Information

### Contributions

W.Y. and T.K. conceived experiments. W.Y. performed imaging experiments and data analysis.

W.Y. and C.M.S. performed stereotaxic surgeries. W.Y., C.M.S., and N.R.R.S. assisted with CRF deletion behavioral experiments. G.A.M. helped with CRF deletion histology, confocal imaging, and quantification. W.Y. and T.K. wrote the paper, with editing contributions from all authors.

### Ethics Declarations

The authors declare no competing interests.

### Additional Information

N/A

